# A critique of the hypothesis that CA repeats are primary targets of neuronal MeCP2

**DOI:** 10.1101/2022.04.26.489598

**Authors:** Kashyap Chhatbar, John Connelly, Shaun Webb, Skirmantas Kriaucionis, Adrian Bird

## Abstract

The DNA binding protein MeCP2 is reported to bind methylated cytosine in CG and CA motifs in genomic DNA, but it was recently proposed that arrays of tandemly repeated CA containing either methylated or hydroxymethylated cytosine are the primary targets for MeCP2 binding and function. Here we investigated the predictions of this hypothesis using a range of published datasets. We failed to detect enrichment of cytosine modification at genomic CA repeat arrays in mouse brain regions and found no evidence for preferential MeCP2 binding at CA repeats. Moreover, we did not observe a correlation between the CA repeat density near genes and their degree of transcriptional deregulation when MeCP2 was absent. Our results do not provide support for the hypothesis that CA repeats are key mediators of MeCP2 function. Instead, we found that CA repeats are subject to CAC methylation to a degree that is typical of the surrounding genome and contribute modestly to MeCP2-mediated modulation of gene expression in accordance with their content of this canonical target motif.

## Introduction

MeCP2 is a chromatin protein that is abundant in neurons and essential for brain function. Initially identified through its affinity for 5-methylcytosine in DNA, the exact nature of its DNA targets has periodically been debated and challenged. Based on in vivo and in vitro data from several laboratories, targeting of MeCP2 to DNA depends on the presence of mC in two motifs: mCG and mCA (the latter mostly in the trinucleotide mCAC) [1–7]. Independent studies suggest that the presence of mCA in neurons is essential, in part at least because of its affinity for MeCP2 [2,8,9]. For example, mice expressing a modified form of MeCP2 that does not interact with mCA but can still bind mCG develop severe Rett syndrome-like phenotypes [8]. In vitro, MeCP2 also binds to the hydroxymethylated (hm) motif hmCAC [3], but the significance of this interaction has been unclear due to the apparent rarity of this modified trinucleotide in brain genomes [10].

This scenario is challenged by a recent proposal that arrayed tandem repeats of the dinucleotide CA are critical MeCP2 targets, exceeding in importance both mCG and isolated mCA moieties as mediators of MeCP2 function [11]. As [CA]_n_ repeat blocks are relatively frequent, it is argued that their proximity to genes provides high affinity ‘landing pads’ through which MeCP2-dependent gene regulation is mediated. Mechanistically, it is suggested that MeCP2 binding to occasional mCA or hmCA moieties within [CA] _n_ repeats can seed cooperative MeCP2 binding across the entire array, including non-methylated CA motifs. Here, we investigate further the relationship between MeCP2 and CA repeats. Our findings do not offer support for the claim that cytosine modification is enriched at [CA]_n_ arrays or that MeCP2 is preferentially bound at these repeat blocks in brain cell nuclei. Moreover, we find that the effects of MeCP2-deficiency on transcription in various brain regions do not correlate with proximity to [CA]_n_ repeats, but instead strongly correlate with local mCAC frequency.

## Results

### Absence of enrichment of modified cytosine in CA repeats

The mouse genome (version mm9) contains ~320,000 [CA]_n_ arrays (minimum 10 base pairs) of variable length with an average of 25 CA repeats each. It has been reported that CA is more frequently methylated or hydroxymethylated within [CA]_n_ repeat arrays than elsewhere in the genome [11], but this comparison did not take account of the preferential methylation of CAC trinucleotides in neurons [3]. While CAC is necessarily very abundant within [CA]_n_ repeats, isolated CA motifs elsewhere in the genome may or may not have C in the third position. Recognising that the trinucleotide sequence CAC is the preferred target of non-CG methylation in neurons and is also a target for MeCP2 binding [3,12], we determine that approximately 6% of all CAC motifs in mouse are found within [CA]_n_ repeats (Figure 1A). Interestingly, humans (version chm13-v1.1) possess a much lower proportion of [CA]_n_ repeats: ~55,000 [CA]_n_ arrays amounting to only ~1% of all CAC motifs in the genome (Figure 1A). Using published data for three mouse brain regions, we confirmed that mCA occurs at a lower frequency outside [CA]_n_, but the frequency of mCAC for each brain region was similar within and outside the repeat arrays (Figure 1B). To determine whether the levels of cytosine modification in [CA]_n_ arrays match the level of mCAC nearby, we plotted mCAC number per gene against mCA number exclusively within CA repeats (Figure 1C). The results showed a strong correlation, indicating that the local density of mCAC in genes is similar regardless of whether the trinuclotide is isolated or within a CA repeat array. We conclude from these findings that, while [CA]_n_ arrays are subject to CAC methylation, they are not targeted preferentially compared with the surrounding genome but tend to adopt a level of mCAC that reflects the surrounding DNA.

**Figure 1.**
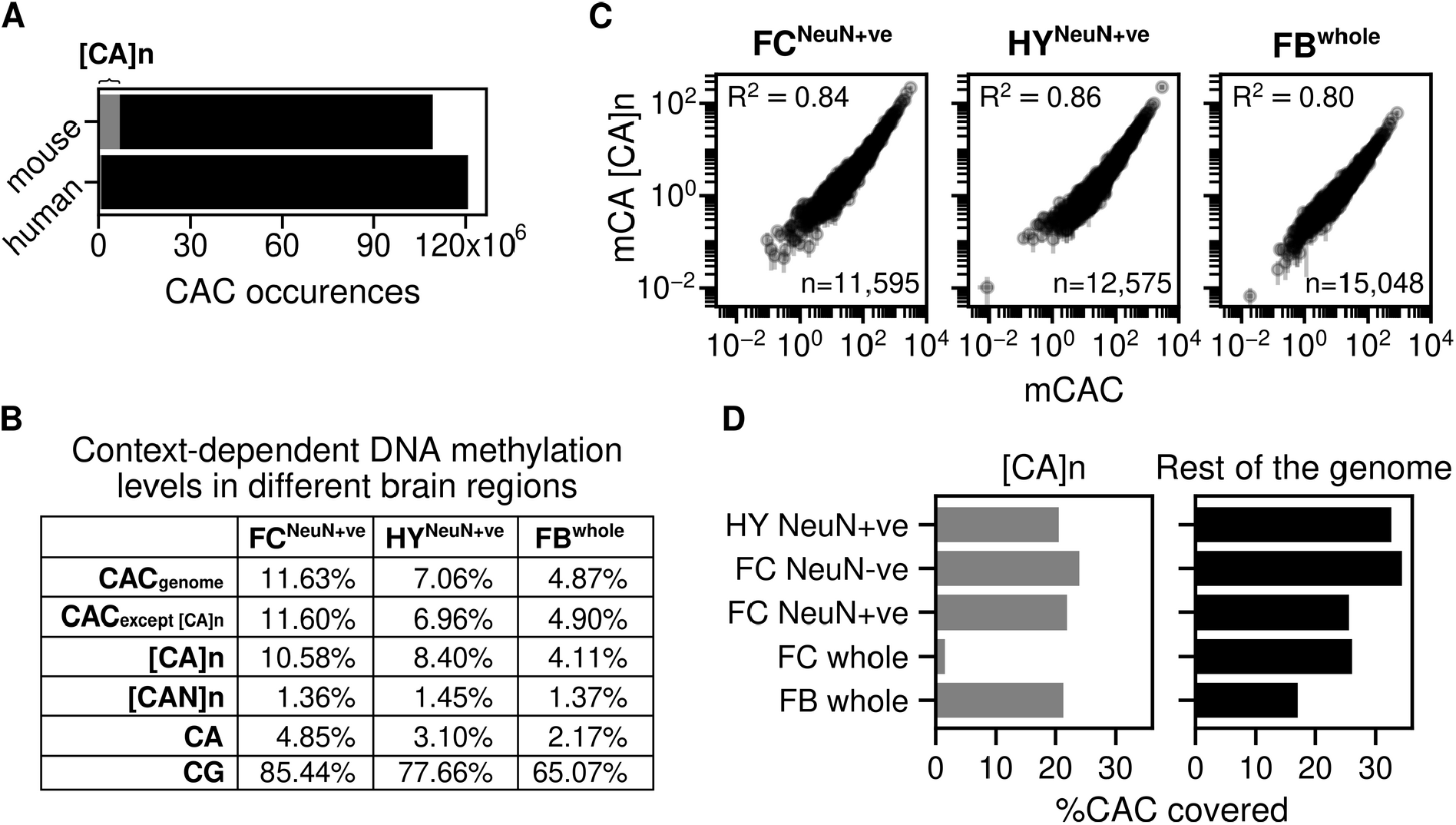
Absence of enrichment of modified cytosines in CA repeat arrays. **A**) Number of CAC occurrences across the mouse and human genome. The grey area corresponds to the number of CAC occurrences within CA repeat arrays. **B**) Genome wide average DNA methylation levels for different cytosine contexts: CAC across the whole genome; CAC across the whole gene except [CA]_n_; [CA]_n_ dinucleotide repeats; [CAN]n trinucleotide repeats (where N is A, T, or G); CA; and CG. DNA methylation levels were quantified from three mouse brain regions: sorted NeuN+ve nuclei from hypothalamus (HY) [3]; sorted NeuN+ve nuclei from frontal cortex (FC) [10] and forebrain (FB) [14]. **C**) mCAC number per gene plotted against mCA number per gene exclusively in CA repeats and Pearson correlation (R2) is calculated for three mouse brain regions: sorted NeuN+ve nuclei from hypothalamus (HY) [3]; sorted NeuN+ve nuclei from frontal cortex (FC) [10] and forebrain (FB) [14]. **D**) Percentage of CAC sites with adequate bisulfite sequence coverage within CA repeat loci and across the mouse genome. The coverage threshold for sorted NeuN+ve Hypothalamus (HY) [3] and whole forebrain (FB) [14] is at least 5 reads and threshold for sorted NeuN+ve, NeuN-ve and whole frontal cortex (FC) [10] is at least 10 reads.

This conclusion assumes that the level of cytosine modification in [CA]_n_ arrays has not been systematically underestimated by bisulfite sequencing due to reduced coverage of repetitive sequences. To test for under-representation in published bisulfite sequence data, we compared the fraction of CAC covered in [CA]_n_ repeats versus the rest of the genome. The results show little difference for forebrain and reduced coverage of [CA]_n_ arrays (<2-fold) in NeuN+ve frontal cortex and hypothalamus (Figure 1D). Slightly lower coverage of CA repeats has little effect on estimates of DNA methylation levels as, even when inadequately covered sequences are excluded, the number of CACs that are reliably detected (1,913,430) is more than sufficient to allow accurate determination of their modification status. The exception to this was the whole frontal cortex dataset (280,015 CACs reliably detected), where bisulfite sequence data gave much lower coverage of [CA]_n_ repeats [10]. Importantly, only this sparsely covered dataset was analyzed by Ibrahim et al (2021). High bisulfite sequence coverage was obtained with purified cortical NeuN+ve (neuronal) and NeuN-ve (mostly non-neuronal) nuclei from the same study [10] (Figure 1D), demonstrating that CA repeats are not intrinsically under-represented in cortex by this technology. Leaving aside the outlier dataset from whole frontal cortex, the evidence indicates a modest bias by bisulfite sequencing against [CA]_n_ repeats. Despite this effect, CA repeat coverage is sufficient to strongly support the conclusion that methylation of mCAC is similar between CA repeats and the rest of the genome.

### Absence of enrichment of MeCP2 binding at CA repeats in brain

We next asked whether MeCP2 is preferentially associated with CA repeat arrays [11]. MeCP2 ChIP of mouse brain reproducibly reveals relatively uniform genome occupancy with few prominent peaks [2,3,14]. This has been interpreted to reflect the high frequency throughout the neuronal genome of short MeCP2 target sites, mCG and mCAC [3]. In contrast, Ibrahim et al report that MeCP2 ChIP-seq reads in cultured mouse embryo fibroblasts (MEFs) are concentrated in prominent peaks coincident with CA repeat clusters [11]. Given the importance of MeCP2 function and the exceptionally high abundance of hmC and mC in neurons, equivalent peaks at [CA]_n_ might be expected in the brain. However, published ChIP data does not support this prediction. As an example, the MeCP2 binding profile (normalised to the KO or input ChIP profiles) across the same ~120 kb region of the mouse genome that was illustrated for MEFs [11] failed to highlight [CA]_n_ arrays (Figure 2A). In view of uncertainties regarding the initial MEF data (see Discussion), our findings question the evidence for preferential localisation of MeCP2 to CA repeats.

**Figure 2.**
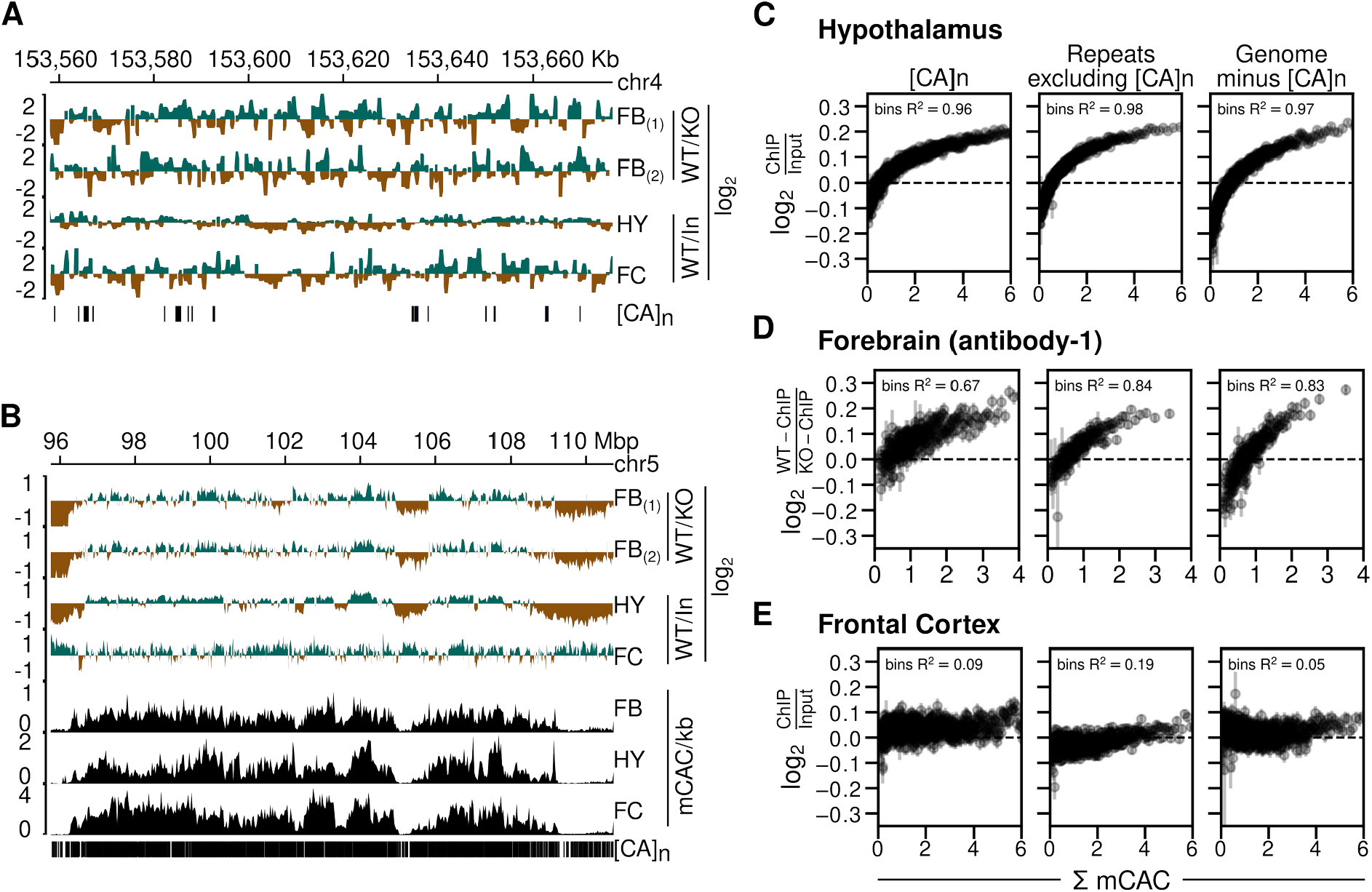
Absence of enhanced MeCP2 binding at CA repeat arrays in brain. **A)** Genome browser screenshots of mouse chr4:153558029-153676791 (version mm9) showing Mecp2 WT ChIP signal normalized to Mecp2 KO ChIP signal for forebrain (FB) using two distinct antibodies [14], Mecp2 WT ChIP signal normalized to input chromatin for hypothalamus (HY) [13] and frontal cortex (FC) [2]. Vertical strokes (bottom row) show location of CA repeat arrays. **B**) As described for panel A but using coordinates at chr5:95751179-110732516. Additionally, DNA methylation tracks showing mCAC/kb for sorted NeuN+ve nuclei from forebrain (FB) [10], hypothalamus (HY) [3] and sorted NeuN+ve nuclei from frontal cortex (FC) [10]. **C**) MeCP2 ChIP signal normalized to input chromatin in mouse hypothalamus [13] plotted against bins of increasing levels of DNA methylation at CAC [3] (shown as mean±standard error of mean) within three different genomic sequence categories. Panels from left to right represent CA repeat loci, simple repeat loci other than CA repeats as identified by RepeatMasker, and 500,000 randomly chosen 1kb genomic windows which do not overlap with CA repeats. R2 values indicate squared spearman correlation from binned mean values of MeCP2 ChIP enrichment and binned mean values of mCAC. **D**) As described for panel C but using data for mouse forebrain [14]. **E**) As described for panel C but using data for mouse frontal cortex [2] and [10].

We also visualised enrichment of MeCP2 at lower resolution across a much larger (15 megabase) region of mouse chromosome 5 (Figure 2B). The results agree with previous reports of a fluctuating distribution of MeCP2 across the genome that broadly tracks the density of mCAC [3]. Although [CA]_n_ repeat arrays are not apparent as prominent sites of MeCP2 binding, their high density at this resolution makes it difficult to discern by inspection alone whether they are preferred. In addition, the global distribution of bound MeCP2 across the neuronal genome limits the value of traditional peak analysis methods to define the binding pattern [3]. To reveal the relationship of ChIP signal to CA repeats versus other genomic regions, we plotted bins of log2-fold change in MeCP2 binding (normalised to KO or input) versus mCAC frequency for three DNA sequence categories: i) [CA]_n_; ii) repeated sequences excluding [CA]_n_; and iii) the rest of genome excluding [CA]_n_. We drew upon published datasets derived from brain regions for which matching ChIP and bisulfite data were available and again normalised to KO ChIP signal or input [3,10,13,14]. The results showed that for hypothalamus, MeCP2 occupancy clearly rose as mCAC frequency increased (Figure 2C). If [CA]_n_ arrays were preferential targets for MeCP2 binding, we would expect them to show elevated ChIP enrichment compared with the other genome categories, but the three DNA sequence types showed closely similar profiles in each brain region. The same conclusion can be drawn from data from forebrain, which is represented by two high-coverage datasets using independent anti-MeCP2 antibodies (Figure 2D). A fourth dataset derived from mouse frontal cortex [2,10] also showed a trio of near-identical ChIP plots, but in this case the relationship to mCAC frequency was less striking (Figure 2E). Rank correlations derived from the averaged bins across the whole genome excluding CA repeats gave R2 values of 0.97 for hypothalamus and 0.83 for forebrain indicating strong dependence between MeCP2 enrichment and mCAC. In the case of frontal cortex an R2 value of 0.05 revealed only modest enrichment for highly methylated regions of the genome. Using peak enrichment to indicate the relative coverage of ChIP-seq datasets, we could identify 271,334 and 236,330 MeCP2 enriched regions in Hypothalamus [13] and Forebrain [14] ChIP data sets respectively, whereas the frontal cortex data [2] detected only 37,817 enriched regions. This ~7-fold decrease suggests lower resolution of the frontal cortex ChIP data which may contribute to its different profile (Figure 2B) and modest correlation between MeCP2 binding and DNA methylation (Figure 2E). Regardless of differences, all three datasets failed to reveal evidence of enhanced binding of MeCP2 at [CA]_n_, as the intensity of ChIP signal at [CA]_n_ was approximately equivalent to that in other parts of the genome.

Based on electrophoretic mobility shift assays, it was further proposed that cooperative binding across [CA]_n_ arrays is facilitated by the affinity of MeCP2 for non-methylated [CA]_n_ [11]. This recalls early reports [15] that certain truncated variants of MeCP2 can bind to CA/TG-rich probes in vitro, although recent evidence failed to validate this mode of binding with full-length MeCP2 either in vitro or in vivo [6]. Ibrahim et al. reported MeCP2 binding to longer non-methylated CA repeat tracts – specifically [CA]_7_. Using a pulldown assay for native MeCP2 binding in mouse brain extracts, however, we failed to detect enhanced MeCP2 binding to [CA]_7_ compared with control DNA probes that lacked CA repeats or in which CA was part of a CAGA repeat array (Figure 3A & B). Introduction of one mC residue into the [CA]_7_ tract significantly increased MeCP2 binding. Moreover, binding displayed a trend towards further enhancement when three more cytosines in the array were methylated. This result is compatible with a linear rather than a cooperative relationship between the amount of mCAC and MeCP2 binding. We noted that 10 mCACs in a non-repetitive probe did not appear to further enhance binding compared with 4 mCACs. Potential explanations for this apparent plateau include steric interference of closely proximal mCAC sites, probe length, etc. Unfortunately, the variability between experiments prevented us from exploring these alternatives quantitatively. Overall, our results fail to confirm an intrinsic affinity of MeCP2 for non-methylated [CA]_7_ in vitro and they suggest that addition of one mCAC motif is not sufficient to cause cooperative MeCP2 binding across a [CA]_7_ array, as further methylation of the [CA]_7_ probe further enhances binding.

**Figure 3.**
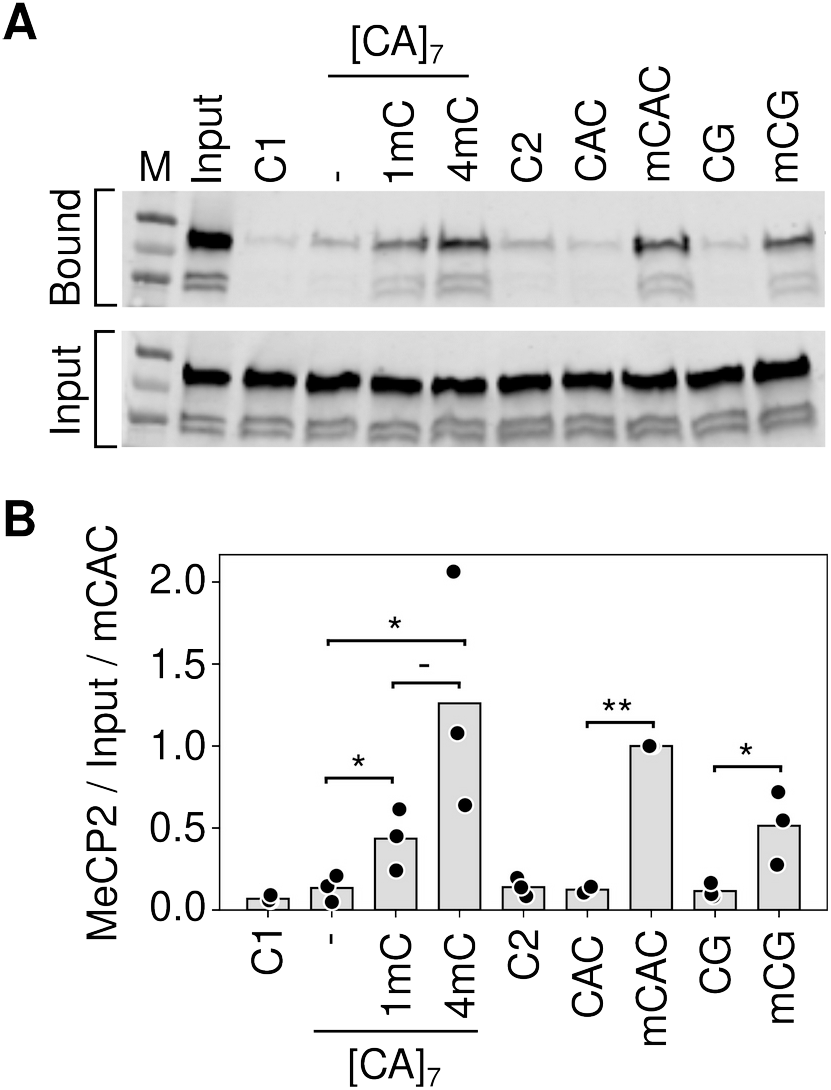
MeCP2 binding to [CA]_7_ is dependent on cytosine methylation. **A)** Example of a pulldown assay for MeCP2 binding using biotin tagged double stranded DNA oligonucleotides incubated with mouse brain nuclear extracts (see Materials and Methods). Unrelated probes C1 and C2 contained no mC. Probe CAC contained 10 non-methylated CACs on one strand, all methylated in mCAC. Alternative versions of the [CA]_7_ probe contained 0, 1 and 4 mCAC motifs labelled as -, 1mC and 4mC respectively. **B**) Quantification of the triplicate data exemplified in A. Significance was estimated using a paired T-test (* pval < 0.05, ** pval < 0.01).

### Minor effect of CA repeats on MeCP2-mediated gene regulation

We next tested the relationship between gene expression changes in the MeCP2-deficient brain and the frequency of [CA]_n_ repeat clusters within gene bodies, drawing on data from independent studies of mouse hypothalamus, forebrain and cortex [2,13,14]. To investigate the effect of enriched [CA]_n_ repeat clusters on transcription, we asked whether significantly up- or down-regulated gene bodies were enriched for [CA]_n_. Box plots of these gene categories failed to show obvious relationship between % [CA]_n_ in up- or down-regulated genes compared to random non-regulated gene bodies (Figure S1A). We also plotted the percentage of all transcription unit nucleotides that belong to [CA]_n_ arrays against the fold change in gene expression when MeCP2 is absent. This differs from a previous analysis [11] by taking into account the direction of transcriptional change and testing multiple brain datasets. Again, the results showed no obvious correlation between the differing levels of [CA]_n_ in the gene body and changes in gene expression in these brain regions (Figure 4, left panels). In contrast, we found that the number of mCAC motifs per gene, either including or excluding [CA]_n_ repeat clusters, correlates positively with the average magnitude of gene up-regulation in the mutant brain (Figure 4, middle panels), supporting the notion that MeCP2 binding to this methylated motif restrains gene expression and confirming previous findings [3,16]. Although gene length correlates with gene misregulation in MeCP2 KO [2], a positive correlation with mCAC persisted when mCAC motifs per gene were normalized to gene length (Figure S1B). This suggests that gene length does sufficient not explain the positive correlation with mCAC in the absence of MeCP2. The strong relationship influenced by mCAC motifs, which is unaffected by inclusion or exclusion of [CA]_n_ arrays, is not expected if CA repeats were the primary drivers of MeCP2-mediated gene regulation. Since CA repeats are subject to DNA methylation at the same level as dispersed CAC motifs (see Figure 1), we expected that the presence mCAC within [CA]_n_ tracts would correlate with gene expression. This was confirmed when the number of mCA motifs in [CA]_n_ repeat blocks was plotted against the fold change in transcription between Mecp2 KO and wildtype brain regions (Figure 4, right panels). To quantify these findings, we calculated the rank correlation with log2 fold change in gene expression for unbinned values. R2 values were consistently higher in plots of total mCAC number per gene (including or excluding [CA]_n_) than for % [CA]_n_ (Figure 4, middle panels). When the level of CA methylation in [CA]_n_ was taken into account (Figure 4, right panels), the correlation with transcriptional change did not exceed that of mCAC elsewhere in the genome. Our findings do not support the hypothesis that cytosine modification in [CA]_n_ has a heightened impact on gene regulation.

**Figure 4.**
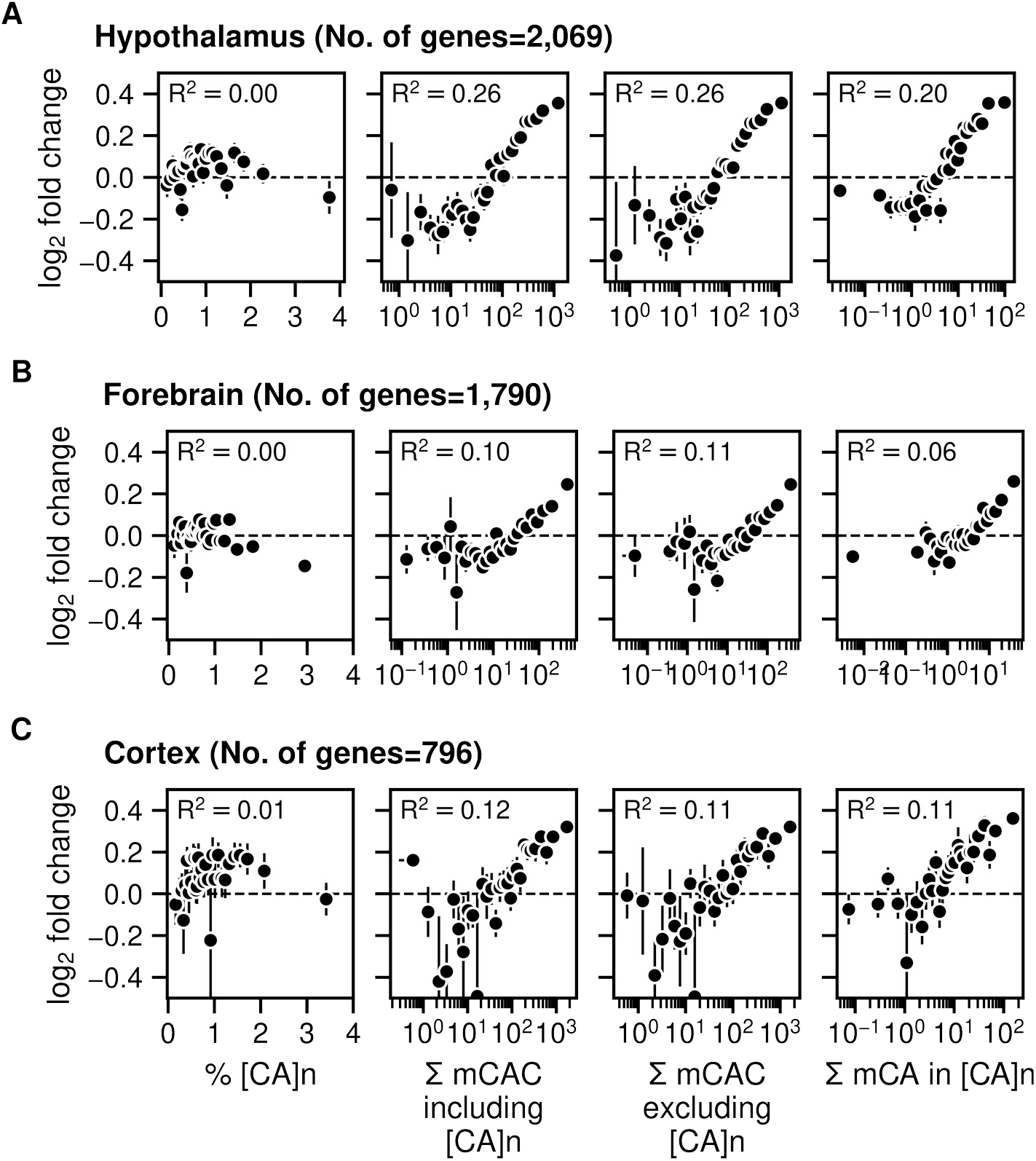
Gene expression increases in Mecp2 KO brain regions according to levels of mCAC but does not correlate with the presence of CA repeat arrays. **A**) Mean log2 fold-change in gene expression in Mecp2 KO versus Mecp2 WT in mouse hypothalamus (differentially regulated genes were defined by padj<0.05) [14] plotted against the percentage of CA repeats within the gene body (left panels); bins of increasing mCAC number per gene including or excluding CA repeats [3] (centre panels); and bins of increasing mCA per gene exclusively within CA repeats [3] (right panels). Spearman rank correlation (R2) is calculated using unbinned values for every panel. **B**) As described for panel A above but using data for mouse forebrain (differentially regulated genes were defined by padj<0.05) [13]. **C**) As described for panel A above but using data for mouse frontal cortex (differentially regulated genes were defined by p-value<0.05) [2,10].

## Discussion

We investigated the possibility that CA repeats are preferred targets of MeCP2 binding due to enrichment of mC or hmC, potentially leading to cooperative binding across the entire tract. In the mouse brain, we detected neither enrichment of CAC modification nor obvious accumulation of bound MeCP2 within [CA]_n_ repeat clusters. Instead, the data derived from several independent studies indicate that while arrays of [CA]_n_ do acquire cytosine methylation, the average level of mCAC within them is typical of the surrounding genome. In agreement with this finding, the level of MeCP2 binding within [CA]_n_ repeats is as expected given the frequency of its target motif mCAC. In the absence of MeCP2, gene expression is up-regulated according to mCAC density as reported previously [1,3,5,13,14,16], but we found no obvious correlation with the proportion of gene bodies made up of [CA]_n_ repeat blocks unless the frequency of mCAC was taken into account. We conclude that the effect of [CA]_n_ tracts on gene expression depends on the amount of mCAC that they contain.

Several further considerations lead us to question the proposed link between hmC and MeCP2 [11]. A major reservation concerns the use of antibody detection of hmC, which provided the initial stimulus for the hypothesis of Ibrahim and colleagues [11]. In their experiments, immunoprecipitation with an antibody directed against hmC revealed prominent apparent peaks of hmC coincident with CA repeat clusters in MEFs. This result is unexpected, as levels of hmC are usually very low in dividing cultured cells compared with neurons. More importantly, others have reported “serious flaws” in the MeDIP method that lead to erroneous reporting of hmC even when both mC and hmC are known to be absent [17]. Strikingly, the resulting false positives, which account for 50-99% of regions identified as “enriched” for DNA modifications, are predominantly found at unmodified short repeat arrays, in particular [CA]_n_. In view of this potentially serious caveat, the evidence for hmC at CA repeats in this MEF cell line must be considered provisional, pending independent biochemical validation.

A second concern relates to the biochemical evidence for the binding specificity of MeCP2. In support of the hypothesis that hmC or mC in CA repeat arrays are the primary targets of MeCP2, Ibrahim et al report that (hmCA)n repeats have a 7-fold higher affinity for MeCP2 in vitro than for the canonical MeCP2 target motif mCG [11]. However, rather than using symmetrically methylated mCG/mCG, which is a validated target sequence, as a comparator, the authors chose hemi-methylated mCG/CG. The affinity of MeCP2 for hemi-methylated mCG/CG is reproducibly little more than background [18–20], making this an invalid control. Estimated dissociation constants for the interaction between MeCP2 and symmetrical mCG/mCG are somewhat variable in the literature depending on the details of the assay, ranging from a low of 400nM [20] to 15nM [18] or 10nM [19]. Notably, these published affinities for mCG/mCG are similar to or higher than the affinity for hmCA-containing repeats reported by Ibrahim et al (410nM).

Finally, the extreme rarity of hmCA in brain is difficult to reconcile with its hypothetical pivotal role. A limitation of many brain methylome datasets is that only bisulfite sequence analysis was performed and therefore mC and hmC are not distinguished. It is clear, however, that while the abundance of neuronal mCA is similar that of mCG in brain, the vast majority of neuronal hmC in confined to hmCG [10]. For example, using a non-destructive method for hmC detection, it was shown that 97.5% of hmC in excitatory neurons is in hmCG, with less than 2.5% in hmCA [21]. This is presumably attributable to the strong preference of Tet enzymes for mCG over mCA as a substrate for mC oxidation [22,23]. While it is possible that hmCA plays roles in gene regulation as suggested in cerebellum [24], it is challenging to deconvolve its roles due to our inability to exclusively eliminate this modification from the genome. We nevertheless consider that the moderate affinity of MeCP2 for this ultra-rare motif offers an unlikely basis for comprehensive new models of MeCP2 function.

## Materials and Methods

### Bioinformatic analyses

#### Sequencing data sets

Table 1 below details the published data sets used for the analyses. These include Chromatin Immunoprecipitation followed by Sequencing (ChIP-Seq), RNA sequencing (RNA-Seq) and Bisulfite sequencing (WGBS-Seq) libraries from different regions of mouse brain quantifying MeCP2 occupancy, gene expression and DNA methylation levels respectively.

**Table 1.** Published data sets used for the analyses.

| Brain tissue | Data set | GEO accession |
| --- | --- | --- |
| Hypothalamus | ChIP-seq | GSE66868 [13] |
| Hypothalamus | WGBS-seq | GSE84533 [3] |
| Hypothalamus | RNA-seq | GSE66870 [13] |
| Forebrain | ChIP-seq | GSE139509 [14] |
| Forebrain | WGBS-seq | GSE128172 [14] |
| Forebrain | RNA-seq | GSE128178 [14] |
| Frontal Cortex | ChIP-seq | GSE67293 [2] |
| Frontal Cortex | WGBS-seq | GSE47966 [10] |
| Visual Cortex | RNA-seq | GSE67294 [2] |

**Table 2.**
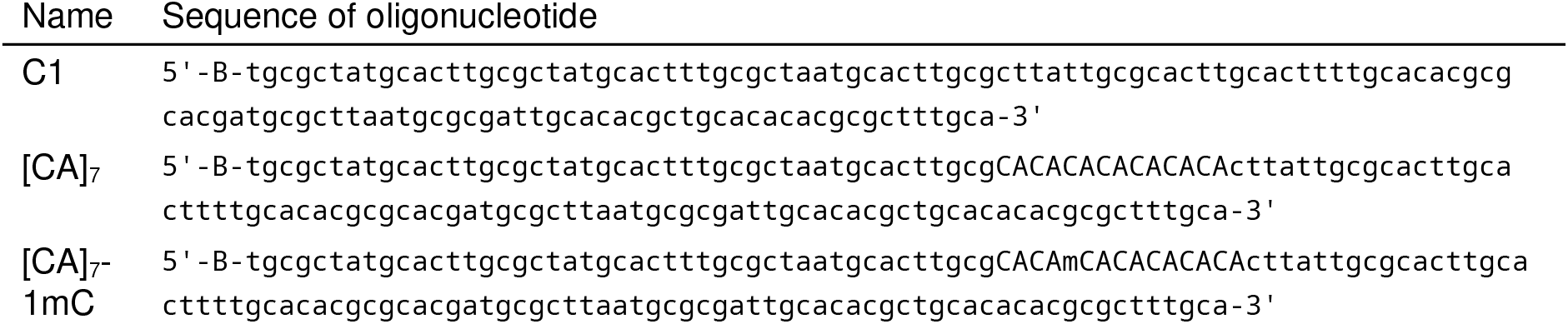
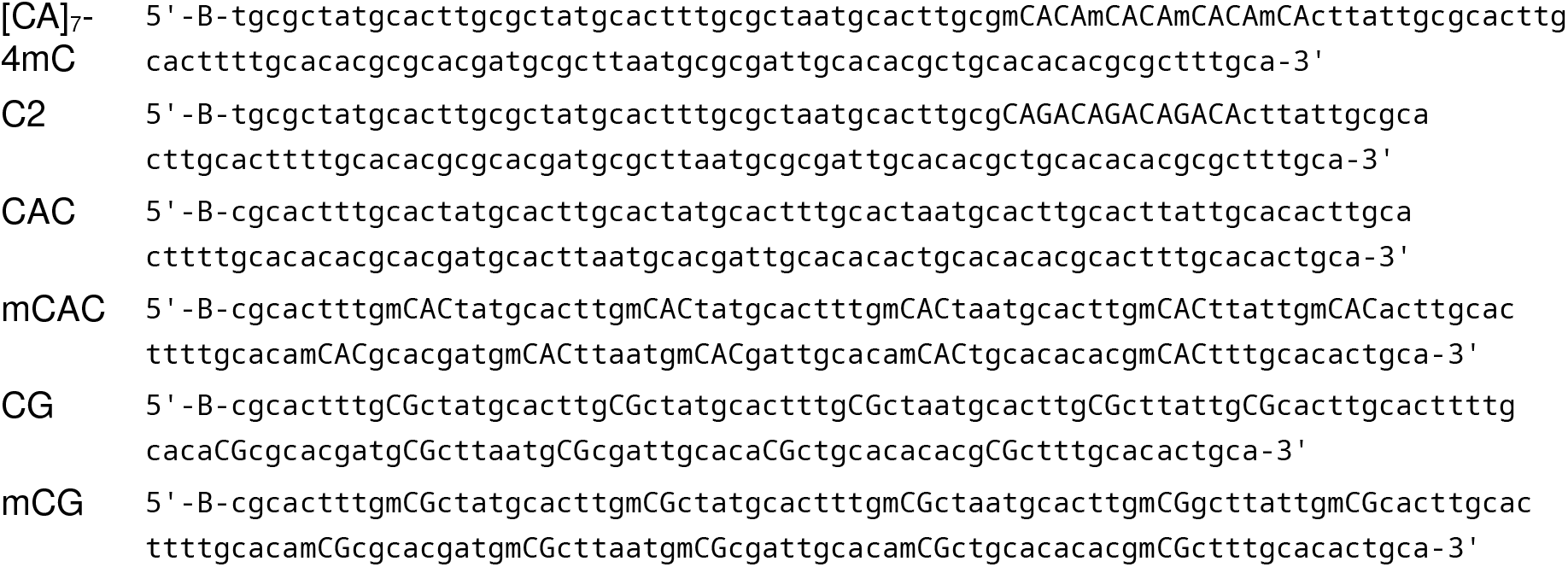
Oligonucleotide sequences for the probes used in the pulldown assay. B=biotin, m=methyl group. All molecules were annealed to the appropriate methylated or non-methylated reverse oligonucleotide.

#### RNA-seq analyses

Gene expression analyses were performed for mouse brain RNA-seq data sets. Raw data were downloaded, mRNA expression was quantified using kallisto [25] and differential expression analysis was performed using DESeq2 [26]. Differentially regulated genes (significance threshold after Benjamini-Hochberg correction p-adjusted value < 0.05 or p-value < 0.05) in MeCP2 KO and WT mouse brain tissue were sorted according to the total amount of mCAC per gene body including [CA]_n_ or excluding [CA]_n_; total amount of mCA in [CA]_n_ of gene body, binned into 30 equal-sized bins and mean log2 fold change of each bin is plotted. Error bars represent the standard error of the mean for that bin. Alternatively, the mean log2 fold change was plotted for differentially regulated genes sorted according to the % CA repeats within the gene body and total methylation within the CA repeats.

#### ChIP-seq analyses

Raw fastq reads were downloaded from GEO and subsequent ChIP-seq analysis was performed on mouse genome (mm9) using snakePipes [27] v2.5.3. log2 ChIP-seq signal over Input signal and log2 Wildtype ChIP-seq signal over MeCP2 Knockout ChIP-Seq signal where available are quantified using bigWigAverageOverBed across genomic locations containing [CA]_n_, Repeats excluding [CA]_n_ and rest of the genome.

#### WGBS-seq analyses

Processed WGBS-Seq data sets described in Table 1 were downloaded and DNA-methylation ratios of individual cytosine nucleotides within CAC, CA and CG contexts was determined across both the sense and non-sense strands. The whole genome is divided into 1 kilobase (kb) windows using bedtools and DNA methylation is calculated in the 1 kb window labelled as mCAC/kb in Figure 2B. For repetitive regions, DNA methylation is calculated within the extended [CA]_n_ or Repeats excluding [CA]_n_ region to match the length of repeat regions equivalent to 1kb. Mean DNA methylation levels for different DNA sequence contexts is calculated for cytosines with adequate sequence coverage. For [CA]_n_ and [CAN]n, all cytosines across both strands within the genomic loci of respective repeats are considered. Because of coverage differences between the bisulfite data sets, we set the cytosine coverage thresholds for hypothalamus [3] and forebrain [14] at 5 compared to more highly covered cortex [10] at at least 10 reads for every cytosine. These thresholds enable reliable estimates of average DNA methylation.

#### [CA]_n_, [CAN]_n_ and Repeats excluding [CA]_n_

The list of genomic locations containing CA and TG repeats was extracted from “Variation and Repeats” group of RepeatMasker track in Mouse (mm9) genome using UCSC table browser functionality [https://genome.ucsc.edu/cgi-bin/hgTables]. For Human (chm13-v1.1), RepeatMasker track was downloaded from processed data [28]. Loci labelled “(CA)n” and “(TG)n” in the RepeatMasker track were used for [CA]_n_. Loci labelled “(CAA)n”, “(TTG)n”, “(CAG)n”, “(CTG)n”, “(CAT)n” and “(ATG)n” were used for [CAN]_n_. Simple repeat sequence loci other than [CA]_n_ are considered as “Repeats excluding [CA]_n_”.

#### CAC occurrences

After extracting the list of genomic loci for [CA]_n_, CAC occurrences are calculated using bedtools and jellyfish for [CA]_n_ and the whole mouse genome.

#### Reproducibility

Source code to reproduce all the analysis and figures is available on the Github repository (https://github.com/kashyapchhatbar/MeCP2_2022_manuscript) and archived at Zenodo (**DOI**: 10.5281/zenodo.6997675).

### Pulldown assay for MeCP2 binding to DNA

This assay was performed as described previously [29] with the following modifications. Biotin-end-labelled double-strand synthetic oligonucleotides (2 μg) were coupled to M280-streptavidin Dynabeads according to manufacturer’s instructions (Invitrogen). Bead-DNA complex was then co-incubated at 4°C for 1.5 h with nuclear protein (10 μg). Nuclear extracts from mouse brain (0.42M salt) were prepared as described [24] and dialysed back into a solution containing 0.15 M NaCl. Following extensive washing, bead-bound proteins were eluted in Laemmli buffer (Sigma) and resolved on a 4–20% SDS-polyacrylamide gel (NEB). The presence of MeCP2 was assayed by western blotting using anti-MeCP2 monoclonal antibody M6818 (Sigma) using IR-dye as a secondary antibody (IRDye 800CW donkey anti-mouse, LI-COR Biosciences). Triplicate assays were scanned then quantified using a LI-COR Odyssey CLx machine and software.

## Acknowledgements

We thank Raphaël Pantier and Matthew Lyst for providing critical feedback on the manuscript. This work was supported by Wellcome Centre grant to A.B., who is also a member of the Simons Initiative for the Developing Brain. K. C. was supported by a scholarship from College of Science and Engineering, University of Edinburgh.

## Author Contributions

Conceptualization, A.B., K.C. and J.C.; Methodology, K.C. and J.C.; Software, K.C.,S.W.; Formal Analysis, K.C.,S.W.; Investigation, K.C., J.C. and S.W.; Writing – Original Draft, K.C., J.C. and A.B.; Writing – Review & Editing, K.C., J.C., S.W., S.K. and A.B.; Supervision, A.B; Funding acquisition, A.B.

## Supplementary figures

**Figure S1.**
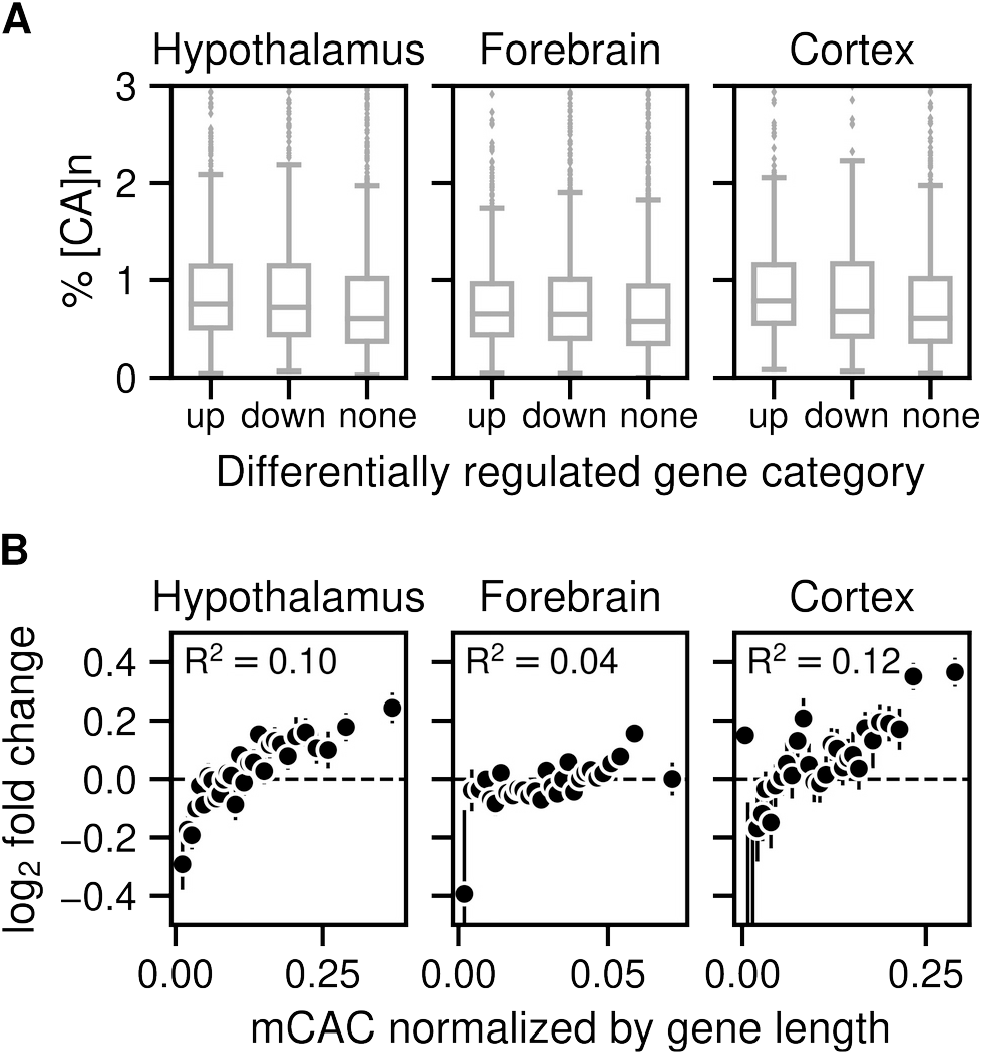
Gene expression increases in Mecp2 KO brain regions according to levels of mCAC but does not correlate with the presence of CA repeat arrays. **A**) Enrichment of [CA]_n_ in up-, down-, or non-regulated genes in Mecp2 KO versus Mecp2 WT for mouse hypothalamus (No. of up-regulated=1,646; down-regulated=1,341 and random subset of non-regulated=2,987 genes) [13]; mouse forebrain (No. of up-regulated=1,359; down-regulated=1,473 and random subset of non-regulated=2,832 genes) [14] and mouse cortex (No. of up-regulated=724; down-regulated=488 and random subset of non-regulated=1,212 genes) [2] **B**) Mean log2 fold-change in gene expression in Mecp2 KO versus Mecp2 WT for mouse hypothalamus [3,13], forebrain [14] mouse cortex [2,10] plotted against the bins of increasing mCAC normalized by gene length. Spearman rank correlation (R2) is calculated using unbinned values for every panel.

## References

1. Guo JU, Su Y, Shin JH, Shin J, Li H, Xie B, et al. Distribution, recognition and regulation of non-CpG methylation in the adult mammalian brain. Nat Neurosci. 2014;17: 215–222. doi:10.1038/nn.3607

2. Gabel HW, Kinde B, Stroud H, Gilbert CS, Harmin DA, Kastan NR, et al. Disruption of DNA-methylation-dependent long gene repression in Rett syndrome. Nature. 2015;522: 89–93. doi:10.1038/nature14319

3. Lagger S, Connelly JC, Schweikert G, Webb S, Selfridge J, Ramsahoye BH, et al. MeCP2 recognizes cytosine methylated tri-nucleotide and di-nucleotide sequences to tune transcription in the mammalian brain. PLOS Genet. 2017;13: e1006793. doi:10.1371/journal.pgen.1006793

4. Lewis JD, Meehan RR, Henzel WJ, Maurer-Fogy I, Jeppesen P, Klein F, et al. Purification, sequence, and cellular localization of a novel chromosomal protein that binds to Methylated DNA. Cell. 1992;69: 905–914. doi:10.1016/0092-8674(92)90610-O

5. Cholewa-Waclaw J, Shah R, Webb S, Chhatbar K, Ramsahoye B, Pusch O, et al. Quantitative modelling predicts the impact of DNA methylation on RNA polymerase II traffic. Proc Natl Acad Sci. 2019;116: 14995–15000. doi:10.1073/pnas.1903549116

6. Connelly JC, Cholewa-Waclaw J, Webb S, Steccanella V, Waclaw B, Bird A. Absence of MeCP2 binding to non-methylated GT-rich sequences in vivo. Nucleic Acids Res. 2020;48: 3542–3552. doi:10.1093/nar/gkaa102

7. Ayata P. Decoding 5HMC as an Active Chromatin Mark in the Brain and its Link to Rett Syndrome. Stud Theses Diss. 2013. Available: https://digitalcommons.rockefeller.edu/student_theses_and_dissertations/238

8. Tillotson R, Cholewa-Waclaw J, Chhatbar K, Connelly JC, Kirschner SA, Webb S, et al. Neuronal non-CG methylation is an essential target for MeCP2 function. Mol Cell. 2021;81: 1260-1275.e12. doi:10.1016/j.molcel.2021.01.011

9. Lavery LA, Ure K, Wan Y-W, Luo C, Trostle AJ, Wang W, et al. Losing Dnmt3a dependent methylation in inhibitory neurons impairs neural function by a mechanism impacting Rett syndrome. West AE, Wassum KM, Zhou Z, editors. eLife. 2020;9: e52981. doi:10.7554/eLife.52981

10. Lister R, Mukamel EA, Nery JR, Urich M, Puddifoot CA, Johnson ND, et al. Global Epigenomic Reconfiguration During Mammalian Brain Development. Science. 2013;341: 1237905. doi:10.1126/science.1237905

11. Ibrahim A, Papin C, Mohideen-Abdul K, Gras SL, Stoll I, Bronner C, et al. MeCP2 is a microsatellite binding protein that protects CA repeats from nucleosome invasion. Science. 2021;372. doi:10.1126/science.abd5581

12. de Mendoza A, Poppe D, Buckberry S, Pflueger J, Albertin CB, Daish T, et al. The emergence of the brain non-CpG methylation system in vertebrates. Nat Ecol Evol. 2021; 1–10. doi:10.1038/s41559-020-01371-2

13. Boxer LD, Renthal W, Greben AW, Whitwam T, Silberfeld A, Stroud H, et al. MeCP2 Represses the Rate of Transcriptional Initiation of Highly Methylated Long Genes. Mol Cell. 2020;77: 294-309.e9. doi:10.1016/j.molcel.2019.10.032

14. Chen L, Chen K, Lavery LA, Baker SA, Shaw CA, Li W, et al. MeCP2 binds to non-CG methylated DNA as neurons mature, influencing transcription and the timing of onset for Rett syndrome. Proc Natl Acad Sci. 2015;112: 5509–5514. doi:10.1073/pnas.1505909112

15. Weitzel JM, Buhrmester H, Strätling WH. Chicken MAR-binding protein ARBP is homologous to rat methyl-CpG-binding protein MeCP2. Mol Cell Biol. 1997;17: 5656–5666. doi:10.1128/MCB.17.9.5656

16. Kinde B, Wu DY, Greenberg ME, Gabel HW. DNA methylation in the gene body influences MeCP2-mediated gene repression. Proc Natl Acad Sci. 2016;113: 15114–15119. doi:10.1073/pnas.1618737114

17. Lentini A, Lagerwall C, Vikingsson S, Mjoseng HK, Douvlataniotis K, Vogt H, et al. A reassessment of DNA-immunoprecipitation-based genomic profiling. Nat Methods. 2018;15: 499–504. doi:10.1038/s41592-018-0038-7

18. Valinluck V, Tsai H-H, Rogstad DK, Burdzy A, Bird A, Sowers LC. Oxidative damage to methyl-CpG sequences inhibits the binding of the methyl-CpG binding domain (MBD) of methyl-CpG binding protein 2 (MeCP2). Nucleic Acids Res. 2004;32: 4100–4108. doi:10.1093/nar/gkh739

19. Hashimoto H, Liu Y, Upadhyay AK, Chang Y, Howerton SB, Vertino PM, et al. Recognition and potential mechanisms for replication and erasure of cytosine hydroxymethylation. Nucleic Acids Res. 2012;40: 4841–4849. doi:10.1093/nar/gks155

20. Lei M, Tempel W, Chen S, Liu K, Min J. Plasticity at the DNA recognition site of the MeCP2 mCG-binding domain. Biochim Biophys Acta BBA -Gene Regul Mech. 2019;1862: 194409. doi:10.1016/j.bbagrm.2019.194409

21. Schutsky EK, DeNizio JE, Hu P, Liu MY, Nabel CS, Fabyanic EB, et al. Nondestructive, base-resolution sequencing of 5-hydroxymethylcytosine using a DNA deaminase. Nat Biotechnol. 2018;36: 1083–1090. doi:10.1038/nbt.4204

22. DeNizio JE, Dow BJ, Serrano JC, Ghanty U, Drohat AC, Kohli RM. TET-TDG Active DNA Demethylation at CpG and Non-CpG Sites. J Mol Biol. 2021;433: 166877. doi:10.1016/j.jmb.2021.166877

23. Hu L, Lu J, Cheng J, Rao Q, Li Z, Hou H, et al. Structural insight into substrate preference for TET-mediated oxidation. Nature. 2015;527: 118–122. doi:10.1038/nature15713

24. Mellén M, Ayata P, Heintz N. 5-hydroxymethylcytosine accumulation in postmitotic neurons results in functional demethylation of expressed genes. Proc Natl Acad Sci. 2017;114: E7812–E7821. doi:10.1073/pnas.1708044114

25. Bray NL, Pimentel H, Melsted P, Pachter L. Near-optimal probabilistic RNA-seq quantification. Nat Biotechnol. 2016;34: 525–527. doi:10.1038/nbt.3519

26. Love MI, Huber W, Anders S. Moderated estimation of fold change and dispersion for RNA-seq data with DESeq2. Genome Biol. 2014;15: 550. doi:10.1186/s13059-014-0550-8

27. Bhardwaj V, Heyne S, Sikora K, Rabbani L, Rauer M, Kilpert F, et al. snakePipes: facilitating flexible, scalable and integrative epigenomic analysis. Bioinformatics. 2019;35: 4757–4759. doi:10.1093/bioinformatics/btz436

28. Hoyt SJ, Storer JM, Hartley GA, Grady PGS, Gershman A, Limouse C, et al. From telomere to telomere: the transcriptional and epigenetic state of human repeat elements analysis code: T2T-CHM13. Zenodo; 2022. doi:10.5281/zenodo.5895031

29. Piccolo FM, Liu Z, Dong P, Hsu C-L, Stoyanova EI, Rao A, et al. MeCP2 nuclear dynamics in live neurons results from low and high affinity chromatin interactions. Dekker J, Zoghbi HY, Dekker J, editors. eLife. 2019;8: e51449. doi:10.7554/eLife.51449

